# FRET-based sensor for measuring adenine nucleotide binding to AMPK

**DOI:** 10.1101/2023.09.05.553069

**Authors:** Roland Abi Nahed, Martin Pelosse, Francesco Aulicino, Florine Cottaz, Imre Berger, Uwe Schlattner

## Abstract

AMP-activated protein kinase (AMPK) has evolved to detect a critical increase in cellular AMP/ATP and ADP/ATP concentration ratios as a signal for limiting energy supply. Such energy stress then leads to AMPK activation and downstream events that maintain cellular energy homeostasis. AMPK activation by AMP, ADP or pharmacological activators involves a conformational switch within the AMPK heterotrimeric complex. We have engineered an AMPK-based sensor, AMPfret, which translates the activating conformational switch into a fluorescence signal, based on increased fluorescence resonance energy transfer (FRET) between donor and acceptor fluorophores. Here we describe how this sensor can be used to analyze direct AMPK activation by small molecules *in vitro* using a fluorimeter, or to estimate changes in the energy state of cells using standard fluorescence or confocal microscopy.

## 1. Introduction

Sensing of available energy within a cell has been an important achievement for the evolution of eukaryotes. It allows rapid responses to fluctuations in energy intake or expenditure, and thus the maintenance of cellular energy homeostasis. The central component within this regulatory framework is the AMP-activated protein kinase (AMPK). This heterotrimeric complex has acquired the capability of being activated by ADP and AMP, while being inhibited by ATP. The affinity for these nucleotides evolved in such a way that the kinase becomes activated by increased ADP/ATP and AMP/ATP ratios (or an adenylate energy charge as defined by Atkinson [1]) just before energy supply becomes limiting for cellular reactions that require free energy [2]. Activated AMPK then relieves such energy stress by triggering a large variety responses specific to cell type and subcellular localization [3]. Most of these either slow down ATP consumption or stimulate ATP regeneration by various downstream mechanisms acting on enzyme activities and gene expression. The major outcome of this enzyme/gene regulation is increased flux through catabolic pathways, mitochondrial proliferation, or even autophagy in extreme situations. Dysregulation of AMPK signaling is involved in various pathologies [4].

Activation of the AMPK heterotrimer by adenine nucleotides (AMP or ADP) or pharmacological compounds involves a conformational switch which we first observed by small angle X-ray scattering [5] and which was later confirmed by electron microscopy and X-ray crystallography [6, 7]. Based on these findings, we engineered the fluorescent sensor AMPfret that translates these conformational changes into a ratiometric fluorescence signal based on fluorescence resonance energy transfer (FRET) [8, 9]. Here, the conformational switch reduces the distance between two fluorophores which increases FRET between the excited donor fluorophore and the acceptor fluorophore. This finally leads to an altered ratio between donor and acceptor emission, the so-called FRET ratio. By this mechanism, AMPfret exploits the naturally evolved sensitivity of AMPK for physiologically relevant cellular ADP/ATP and AMP/ATP ratios, and thus reports allosteric regulation of AMPK. However, AMPfret fluorescence is unchanged when AMPK is covalently activated (phosphorylated at α-T172) by upstream kinases, likely because this does not trigger the required conformational change. AMPfret thus visualizes endogenous energy stress signaling, while entirely preserving AMPK structure, regulation and enzyme activity. Moreover, AMPfret reports binding of pharmacological AMPK activators and is therefore a suitable tool to screen for potential AMPK activators or to dissect non-covalent activation mechanisms of AMPK in general [8].

It is also possible to exploit AMPfret as a reporter system for monitoring energy state at a single cell level in space and time. Several different genetically-encoded fluorescent protein sensors for ATP or ADP/ATP ratios have been generated in recent years. These include A-team [10], Perceval and PercevalHR [11, 12], QUEEN [13], iATPSnFRs [14] or ChemoG–ATP [15]. They represent a major advance for studying cellular energy metabolism, and hold the promise to overcome the limitations of conventional biochemical and low-resolution techniques [16]. However, it is unclear which range of intracellular adenylate ratios are appropriately monitored. These sensors may deliver signals in an adenylate concentration range that is not necessarily physiologically meaningful. AMPfret has the advantage of generating signals from adenylate ratios that represent an energy stress for the cell. It works at the single cell level, without lag-phase and in a fully reversible manner. However, its sensitivity may also limit its use for transient transfection of cell cultures, since transfection protocols already induce an energy stress. This can restrict the dynamic range of the FRET ratio that remains exploitable for subsequent experiments. To overcome this, cell lines can be used that stably express AMPfret at appropriate levels.

Here we present vectors and applications for all these different approaches. This includes bacterial expression and purification of recombinant AMPfret for *in vitro* fluorometric FRET assays to determine allosteric AMPK activation (e.g. by adenylates or pharmacological activators), as well as transient or stable expression of AMPfret in mammalian cell lines, and their measurement by fluorescence and confocal microscopy.

## 2. Materials

### 2.1. Assembly of vectors for AMPfret expression in bacterial and mammalian cells

1. pACE α2-mseCFPΔ11, pDC β2 and pDS γ1-cp173Venus plasmids for bacterial expression and pACEMam2 α2-mseCFPΔ11, pMDK-β2 and pMDS-γ1-cp173Venus for mammalian expression. Empty backbones are part of the MuliColi and MultiMam expression systems (Geneva Biotech)
2. Phusion high-fidelity polymerase kit
3. T4 DNA polymerase
4. 10 mM dCTP or any other single nucleotide solutions
5. Cre recombinase kit
6. BW23474 chemical competent cells or equivalent for propagation of donor plasmids
7. Top10 chemical competent cells or equivalent for propagation of acceptor and Cre-assembled plasmids
8. Heat block
9. Thermocycler
10. QIAquick Gel Extraction Kit
11. QIAprep Spin Miniprep Kit
12. Agarose gel electrophoresis system
13. Agarose Type D-5 DNA-grade
14. GelRed
15. 50x TAE Buffer: 2 M Tris, 1 M acetic acid, 50 mM EDTA pH 8
16. 1 kb DNA Ladder and 100 bp DNA Ladder
17. LB medium
18. LB-agar
19. Plastic petri dishes
20. Ampicillin: 100 mg/mL in ddH2O (1000x stock solution), store at −20° C
21. Gentamycin: 10 mg/mL in ddH2O (1000x stock solution), store at −20° C
22. Kanamycin: 50 mg/mL in ddH2O (1000x stock solution), store at −20° C
23. Spectinomycin: 50 mg/mL in ddH2O (1000x stock solution), store at −20° C
24. Chloramphenicol: 30 mg/mL in 80% ethanol (1000x stock solution), store at −20° C
25. Bacterial spreaders, single use
26. Sterile hood or Bunsen burner

### 2.2. Assembly of vectors for generating mammalian cell lines with stable AMPfret expression

#### 2.2.1. In silico CRE-mediated recombination

1. Sequences of acceptor and one or more donor plasmids in “Fasta” or “GenBank” format can be found at https://doi.org/10.6084/m9.figshare.21695555.v1
2. CRE-rec ACEMBLER software [17]
3. SnapGene software (Dotmatics, Boston, MA, USA; available at <u>snapgene.com</u>)

#### 2.2.2. In vitro CRE-mediated recombination

1. 0.2 mL PCR tubes
2. CRE recombinase (NEB)
3. 10x CRE-recombinase reaction buffer: 330 mM NaCl, 500 mM Tris-HCl, 100 mM MgCl_2_ (pH 7.5 at 25 °C)
4. One acceptor and one or multiple donor plasmids in equimolar ratios. Total DNA amount should be between 1000 and 2000 ng per reaction.

#### 2.2.3. Transformation into OneShot Top10 E. coli

1. OneShot Top10 chemically competent *E. coli*, store at −80° C (Invitrogen)
2. Autoclaved LB
3. Autoclaved LB agar
4. Sterile Petri dishes
5. Sterile ddH2O
6. Gentamycin 10 mg/mL in ddH2O (1000x stock solution), store at −20° C
7. Kanamycin 50 mg/mL in ddH2O (1000x stock solution), store at −20° C
8. Spectinomycin 50 mg/mL in ddH2O (1000x stock solution), store at −20° C
9. Chloramphenicol 30 mg/mL in 80% ethanol (1000x stock solution), store at −20° C
10. Bacterial spreaders, single use
11. 15 mL Falcon tubes

#### 2.2.4. Miniprep and screening

1. QIAprep Spin Miniprep Kit
2. 0.2 mL PCR tubes
3. Restriction endonucleases: EcoRV, EcoRI, BamHI, HindIII, KpnI/ScaI
4. CutSmart buffer (or appropriate endonuclease buffer, depending on brand)
5. TAE buffer (50x): 2.0 M Tris base, 1.0 M Acetic acid, 0.05 M EDTA pH 8.4 (25 °C)
6. Agarose
7. DNA staining dye
8. Gel casting unit and comb
9. 6x DNA loading dye
10. 1 kb plus DNA ladder (NEB)

### 2.3. Expression of recombinant AMPfret in E. coli for in vitro experiments

1. BL21 DE3 star chemical competent cells (Invitrogen)
2. pACEMBL AMPfret 2.1 (from Section 3.1.1)
3. Autoclaved LB
4. LB-agar plate containing 100 µg/mL ampicillin, 30 µg/mL chloramphenicol and 50 µg/mL spectinomycin
5. Ampicillin 100mg/mL in ddH2O (1000x stock solution), store at −20° C
6. Spectinomycin 50 mg/mL in ddH2O (1000x stock solution), store at −20° C
7. Chloramphenicol 30 mg/mL in 80% ethanol (1000x stock solution), store at −20° C
8. Autoinducing medium (0.5% glycerol, 0.05% glucose, and 0.2% lactose) [18]
9. Rotary shaker incubator with cooling (18-37 °C)
10. OD reader (Biowave 3000 CO8000 Cell Density Meter) and plastic cuvettes
11. Centrifuge, rotor and suitable containers for 6 L of bacterial culture.
12. Liquid N_2_

### 2.4. Purification of recombinant AMPfret for in vitro experiments

1. Akta purifier (or similar liquid chromatography system for proteins)
2. LC injection device for larger volumes (Super-loop or equivalent)
3. Cold room (4°C)

#### 2.4.1. Cell lysis

1. Sonicator
2. Centrifuge and rotor capable of 75 000 g
3. Lysis buffer: 0.5 M sucrose, 30 % glycerol, 50 mM Tris pH 8, 100 mM NaCl, 2 mM MgCl_2_, 2 mM β-mercaptoethanol, 20 mM imidazole and protease inhibitors (Roche)

#### 2.4.2. Immobilized metal affinity chromatography (IMAC)

1. 5 mL Ni-NTA Superflow resin
2. Wash buffer: 50 mM Tris pH8, 100 mM NaCl, 20 mM imidazole, 2 mM MgCl_2_, 2 mM β-mercaptoethanol
3. High salt buffer: wash buffer + 1 M NaCl
4. Elution buffer: wash buffer + 400 mM imidazole

#### 2.4.3. SDS-PAGE of chromatography fractions

1. Electrophoresis chamber
2. Powerpack
3. Mini-Protean TGX gels 4-20 % Precast gels (Tris-glycine, 10 well, 50 µl)
4. Blue loading buffer pack, 3x supplemented with 42 mM DTT
5. Precision Plus Protein™ Dual Color Standards
6. Running buffer: 25 mM Tris, 190 mM glycine, 0.1 % SDS, pH 8.3
7. InstantBlue staining buffer

#### 2.4.4. Heparin affinity chromatography

1. 5 mL Heparin column
2. Buffer A :50 mM Tris pH 8, 100 mM NaCl, 2 mM MgCl_2_, 2 mM β-mercaptoethanol
3. Buffer B: 50 mM Tris pH8, 1M NaCl, 2 mM MgCl_2_, 2 mM β-mercaptoethanol

#### 2.4.5. Size exclusion chromatography (SEC)

1. Amicon™ Ultra-4 /-15 centrifugal filter units (50 kDa)
2. Cooled table top centrifuge
3. Superose 6 10/300 gel filtration column
4. SEC buffer: 50 mM Tris pH 8, 200 mM NaCl, 2 mM MgCl_2_, 2 mM β-mercaptoethanol

#### 2.4.6. Storage

1. Amicon™ Ultra-4 /-15 centrifugal filter units (50 kDa)
2. Device for photometric protein determination (Nanodrop or equivalent)
3. Glycerol anhydrous

### 2.5. In vitro FRET assays using fluorimetry

1. Fluorimeter PTI (Photon Technology International)
2. Felix software for data acquisition (PTI)
3. Quartz cuvette
4. Vortex mixer
5. Spectro buffer: 50 mM Tris pH 8, 200 mM NaCl, 5 mM MgCl_2_, 2 mM β-mercaptoethanol
6. AMPfret 2.1 construct at 3 mg/mL in 50 % glycerol (v/v) (from Section 3.4)
7. 1.5 mM AMP, prepared in Spectro buffer
8. Microsoft excel or SigmaPlot 13.0 for data treatment

### 2.6. Cell-based FRET assays using microscopy

#### 2.6.1. Cell culture and transfection

1. HEK 293T cells
2. FBS
3. Trypsin
4. DMEM 4.5 g/L Glucose
5. Penicillin/streptomycin
6. 8-well chambered coverglass with non-removable wells (e.g. LabTek; ThermoFisher ref. 155409)
7. PBS with calcium and magnesium
8. MultiMam AMPfret 2.1 (from Section 3.1.2)
9. OptiMEM
10. PolyFect transfection reagent
11. Lipofectamine 3000
12. Black 96-well plate
13. Black 12-well plate
14. 6-well plate
15. T75 flask
16. Puromycin
17. Hygromycin

#### 2.6.2. FRET assays with confocal or epifluorescence microscopy

1. Epifluorescence microscope (e.g. Leica DMi8 equipped with a HC PL APO 40x/0.95 dry objective) or confocal microscope (e.g. Leica TCS SP8 equipped with a Plan Apo 63x/1.40 oil objective) or equivalent
2. FRET module software (LAS X FRET SE module from Leica)
3. Incubation chamber for microscope
4. 2-well slide for microscopy (e.g. µ-Slide 2 Well Polymer, IBIDI).
5. DMEM containing 4.5 g/L glucose, 10 % FBS, penicillin and streptomycin
6. 2 mM 5-aminoimidazole-4-carboxamide ribonucleotide (AICAR) in complete medium, 200 µL per well
7. FluoroBrite medium supplemented with 10 % FBS, 1 mM L-glutamine and 1 % Penicillin/Streptomycin
8. HEK 7.01 stable cell line
9. 1 mM FCCP (carbonyl cyanide 4-(trifluoromethoxy)phenylhydrazone) in absolute ethanol.
10. FIJI image processing package (https://imagej.net/software/fiji/)
11. Excel
12. GraphPad Prism (Dotmatics, Boston, MA, USA; available at www.graphpad.com) or similar

### 2.7. SDS-PAGE and immunoblotting

1. PBS
2. Homogenization buffer: RIPA buffer 1x, ready-to-use solution containing 150 mM NaCl, 1.0 % IGEPAL CA-630, 0.5 % sodium deoxycholate, 0.1 % SDS, 50 mM Tris, pH 8.0.
3. EDTA-free antiprotease cocktail
4. Phosphatase inhibitor cocktail
5. Smart micro-BCA protein assay kit
6. Plate reader capable of measuring absorbance at 562 nm.
7. Blue loading buffer pack, 3x supplemented with 42 mM DTT
8. Precision Plus Protein™ Dual Color Standards
9. Electrophoresis chamber
10. Mini-Protean TGX gels 4-20 % Precast gels (tris-glycine, 10 well, 50 µl)
11. Powerpack
12. Running buffer: 25 mM Tris, 190 mM glycine, 0.1 % SDS, pH 8.3
13. Protein transfer equipment
14. Nitrocellulose membrane
15. TBST: 20 mM Tris–base, pH 7.5, 150 mM NaCl, 0.1 % Tween 20
16. Blocking solution: TBST with 5 % skimmed milk
17. Primary antibody: anti-α-AMPK, anti-β-AMPK, anti-γ-AMPK anti-P-Thr172-AMPKα, anti-ACC, anti-P-Ser79-ACC and anti-pan-actin antibodies (all Cell Signaling)
18. Horseradish peroxidase-conjugated antibody solution (Goat anti-rabbit, Jackson): 1:10,000 in blocking solution
19. ECL Select Western Blotting Substrate Kit

## 3. Methods

### 3.1. Assembly of vectors for AMPfret expression in bacterial and mammalian cells

This and the following section provide information on how to generate AMPfret vectors for different applications. Vectors for bacterial expression use the MutliColi expression system (see Section 3.1.1.). They allow generation of recombinant AMPfret for in vitro quantification of nucleotide-induced conformational changes within the AMPK heterotrimer. Vectors for transient transfection of mammalian cells are based on the similar MultiMam expression system (see Section 3.1.2.). They enable AMPfret-based monitoring of adenine nucleotides in living cells, which is limited, however, by energy stress already imposed by the transfection procedure itself. Finally, generation of cell line clones stably expressing all three AMPfret subunits is the most reliable manner to monitor adenine nucleotides in mammalian cells, although clone isolation may be time consuming. One strategy to obtain such cells involves all-in-one vectors for targeted integration (see Section 3.2.).

#### 3.1.1. Vector assembly into MutliColi expression system for bacterial expression

Genes encoding for the AMPK subunits α2 (PRKAA2; Gene ID: 78975), β2(PRKAB2; Gene ID: 5565), γ1 (PRKAG1; Gene ID: 25520) and the mseCFPΔ11/cp173Venus FRET pair, were PCR amplified using Phusion polymerase and assembled by SLIC into pACE1, pDC and pDS of the MultiColi expression system as described in [8]. Subsequent single-subunit coding plasmids (i.e. pACE α2-mseCFPΔ11, pDC β2 and pDS γ1-cp173Venus), verified by restriction digest and sequencing, were assembled via Cre-LoxP recombination to give pACEMBL AMPfret 2.1 as described below and in [19].

1. In 0.2 mL PCR tubes, mix acceptor and donors plasmids in equimolar ratio for a total DNA amount of 1000 ng in ddH2O. Total volume of DNA and water should not exceed 8 μL.
2. Add 1 μL of 10x CRE-recombinase buffer.
3. Add 15 U of CRE recombinase.
4. Add ddH_2_O to 10 µl final volume.
5. Mix by pipetting and spin down.
6. Incubate at 37 °C for 1 h.
7. Stop the reaction by heat inactivation at 70° C for 10 min.

The reaction can be now transformed into chemically competent OneShot Top10 *E. coli* or stored at −20 °C

#### 3.1.2. Vector assembly into MultiMam expression system for transient mammalian cell expression

For transient mammalian cell expression, the same process as in 3.1.1. was followed except AMPK-α2-mseCFPΔ11, -β2 and -γ1-cp173Venus were cloned into pACEMam2, pMDK and pMDS, respectively.

### 3.2. Assembly of vectors for generating mammalian cell lines with stable AMPfret expression

Vectors for MultiMam AMPfret (see Section 3.1.2.) were engineered to be included in an all-in-one vector for large cargo integration at the human ACTB locus using CRISPR/Cas9 technology as previously described [20]. An all-in-one acceptor (pMultiMate HITI-2c 4K-CH) encodes Cas9, ACTB sgRNA and homology independent targeted integration (HITI-2c) donor. When successfully transfected into mammalian cells, pMultiMate HITI-2c 4K-CH produces an intronic double strand break (DSB) upstream of the last ACTB coding exon. The HITI-2c donor is excised from the plasmid backbone by Cas9, and inserted at the genomic DSB site through non-homologous end-joining. The resulting editing outcome is the insertion of a synthetic exon coding for ACTB::T2A::H2BiRFP::P2A::Puromycin-R (5’ integration marker) and CMV Hygromycin-R (3’ integration marker) (**Figure 1a**). A loxP site in between 5’ and 3’ integration markers allows for iterative functionalization of the intervening DNA payload by CRE-mediated recombination *in vitro* (**Figure 1b**).

**Figure 1.**
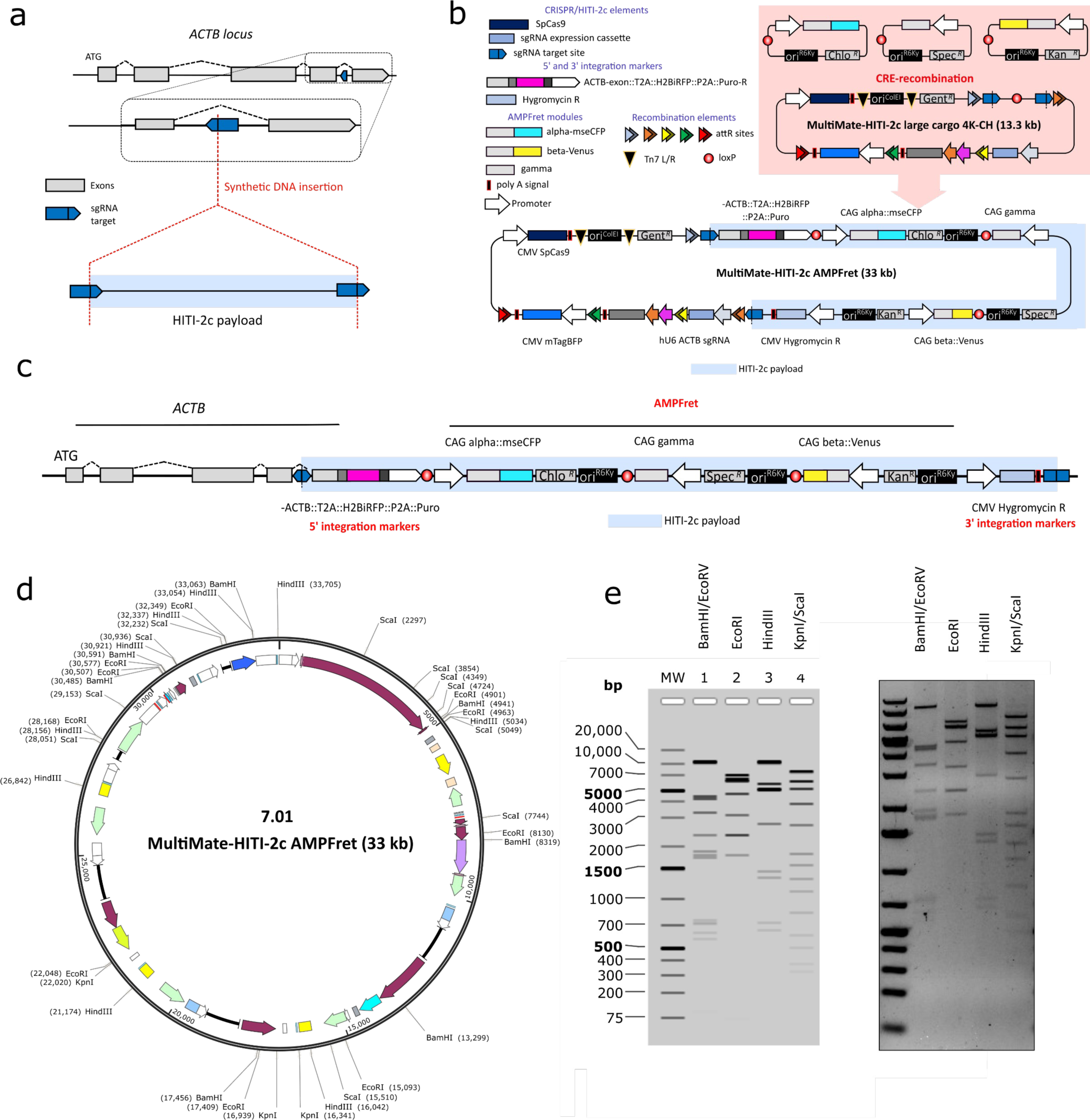

To perform in-vitro CRE-mediated recombination, pMDC α2-mseCFPΔ11 was generated through Gibson cloning by transferring α2-mseCFPΔ11 from pACEMam2 (see Section 3.1.2.) to pMDC and used together with pMDK β2-cp173Venus and pMDS γ1 already generated before (see Section 3.1.2.). The final CRE-recombination reaction consisted of one acceptor (pMultiMate HITI-2c 4K-CH) and three donors (pMDC α2-mseCFPΔ11, pMDK β2-cp173Venus and pMDS γ1) (**Figure 1b**). The resulting vector (7.01) is subsequently used to transfect HEK 293T cells and generating stable cell lines carrying the AMPfret modules integrated at the ACTB locus (**Figure 1c,d**).

#### 3.2.1. In silico CRE-mediated recombination

1. Open CreRec ACEMBLER software.
2. Import sequences for 1 acceptor (MultiMate HITI-2c H2BiRFP PH) and the three AMPFret donors (pMDC alpha2-mseCFP, pMDK beta2-Venus and pMDS gamma1) (found at Figshare, https://doi.org/10.6084/m9.figshare.21695555.v1).
3. Assemble the CRE-reaction by using the “CRE” button.
4. Set the desired ratio for acceptor/donors to 1.
5. Save assembled CRE reactions individually, these will be versions of the same construct differing also by their topology.
6. Import the assembled CRE reactions files in Snapgene and simulate agarose gels after digestion with:

BamHI/EcoRV.
EcoRI,
HindIII
KpnI/ScaI.
(see **Figure 1e**)
7. Save the results.

#### 3.2.2. In vitro CRE-mediated recombination

8. In 0.2 mL PCR tubes, mix acceptor and donors plasmids in equimolar ratio for a total DNA amount of 1000 ng - 2000 ng in ddH_2_O. Total volume of DNA and water should not exceed 8 μL. For equimolar ratio rates, use 501 ng of MultiMate HITI-2c H2BiRFP PH, 180 ng of pMDC alpha2-mseCFP, 167 ng of pMDK beta2-Venus and 150 ng of pMDS gamma1.
9. Add 1 μL of 10x CRE-recombinase buffer.
10. Add 15 U of CRE recombinase.
11. Add ddH_2_O to 10 µl final volume.
12. Mix by pipetting and spin down.
13. Incubate at 37 °C for 1 h.
14. Stop the reaction by heat inactivation at 70° C for 10 min.
15. The reaction can be now transformed into chemically competent OneShot Top10 *E. coli* or stored at −20 °C.

#### 3.2.3. Transformation into OneShot Top10 E. coli

1. Pre-warm autoclaved LB at 37 °C.
2. Thaw one aliquot of OneShot Top10 *E. coli* on ice.
3. Add 5 μL of the CRE reaction and incubate on ice for 20 min.
4. Incubate the vial at 42 °C for 45 s.
5. Quickly transfer the vial on ice and incubate for 2 min.
6. Add 1 mL of warm LB.
7. Incubate the vial for 2 h at 37° C while shaking. For optimal recovery efficiency, place the vial horizontally in a shaking incubator.
8. For each reaction, prepare one Petri dish with 15 mL LB agar and appropriate antibiotics (see below).
9. Heat the autoclaved LB agar in a microwave until completely melted.
10. Allow the LB agar to cool down for 10 min.
11. Near the flame of a Bunsen burner, add all antibiotics in precursor plasmids. In this case: 10 μg/mL Gentamycin, 50 μg/mL Kanamycin, 50 μg/mL Spectinomycin and 30 μg/mL Chloramphenicol), mix by pipetting and transfer 15 mL to each Petri dish.
12. Allow the plate to cool down.
13. Centrifuge the vial of transformed Top10 cells at 3,000 x g for 5 min and discard the supernatant by decanting into a waste vessel.
14. Near the flame of a Bunsen burner, resuspend the bacterial pellet in 200 ul of fresh LB and spread on the Petri dish with LB agar and appropriate antibiotics.
15. Transfer the plate to a static incubator and incubate at 37 °C overnight.
16. Near the flame of a Bunsen burner, pick 3-5 individual colonies using sterile pipette tips and transfer each colony to a 15 mL Falcon tube.
17. Pour 3 mL of LB supplemented with the appropriate antibiotics (see above) into each 15 mL Falcon tube and incubate overnight while shaking at 37 °C.

#### 3.2.4. Miniprep and screening

1. Pellet bacterial culture by centrifugation at 4000 rpm for 5 min.
2. Discard the supernatant and extract plasmid DNA using the Qiagen Miniprep kit following manufacturer’s recommendation.
3. Elute the plasmid DNA in 50 µL elution buffer.
4. Transfer 5 µL of each miniprep to a 0.2 mL PCR tube.
5. For a single restriction digestion, add 2 µL 10x CutSmart buffer and 0.5 µL restriction endonuclease and ddH2O to a final volume of 20 µL. Any single or double restriction digest that provides information on the given construct can be chosen, e.g. those mentioned above in Section 3.2.1. (point 6) and illustrated in **Figure 1e**, or others found in **Figure 1d** or the plasmid maps at Figshare (DOI: 10.6084/m9.figshare.21695555.v1).
6. Spin down briefly and incubate at 37° C for 30 min.
7. Prepare a 1 % agarose gel in 1x TAE, heat in microwave until the agarose is completely dissolved and supplement it with staining dye e.g. GelRed (1:10000).
8. Pour the gel into a gel casting unit and add the appropriate comb, let it cool until solid.
9. Transfer the gel into a gel running unit, pour 1x TAE until completely covered and remove the comb.
10. Load 5 µL 1 kb plus DNA ladder into the first well.
11. Add 4 µL 6x DNA loading dye to each reaction, mix by pipetting and load the whole content of each reaction into a single well.
12. Run the gel until the bromophenol blue/purple dye reaches the bottom.
13. Image using a blue light transilluminator or a gel imager.
14. Compare results with in silico digestion obtained from *in silico* CRE recombination step (**Figure 1e**).
15. Determine plasmid topology and store plasmid DNA at 4 °C short term, or −20 °C for long term storage.

### 3.3. Expression of recombinant AMPfret in E. coli for in vitro experiments

1. Transform 50 µL of BL21 DE3 star chemically competent cells with 3 µL pACEMBL AMPfret 2.1 (from Section 3.1.1). For detailed transformation instructions, see Section 3.2.3.
2. Spread the transformation mixture on an LB-agar plate containing 100 µg/mL ampicillin, 30 µg/mL chloramphenicol and 50 µg/mL spectinomycin.
3. Incubate overnight at 37°C.
4. Pick from the agar plate a single colony and inoculate 200 mL LB containing 100 µg/mL ampicillin, 30 µg/mL chloramphenicol and 50 µg/mL spectinomycin.
5. Let the culture grow overnight at 37 °C in a shaking incubator.
6. Measure the OD_600_ of the overnight culture (typical OD_600_ = 2 to 3).
7. Use the overnight culture to inoculate 6 L of autoinducing medium supplemented with appropriate antibiotics, such that the final OD_600_ = 0.05, and culture at 37 °C.
8. Record culture OD_600_ regularly (every 60 min).
9. When the OD_600_ reaches 0.8, cool the incubator to 18 °C and let the culture grow until the next day.
10. After overnight expression, transfer bacteria into centrifuge containers.
11. Pellet the cells by centrifugation at 4,000 x g for 30 min at 4 °C.
12. Decant supernatant and flash freeze the pellets in liquid nitrogen and store them at −80 °C until further purification.

### 3.4. Purification of recombinant AMPfret for in vitro experiments

All purification steps are carried out at 4 °C or on ice using cold buffers. Never freeze AMPfret protein during the purification procedure. Preserve it in 50 % glycerol at −20 °C to avoid freezing of the sample (see 3.4.6.).

#### 3.4.1. Cell lysis

1. Weight the bacterial pellet and resuspend it in ice cold lysis buffer (1 g pellet / 10 mL lysis buffer).
2. Lyse the bacteria on ice by sonication 5 s ON / 30 s OFF, 70 % power (total of 1 min ON per gram of pellet).
3. Clarify the lysate by centrifugation at 75,000 x g for 45 min at 4 °C

#### 3.4.2. Immobilized metal affinity chromatography (IMAC)

1. Apply the supernatant on 5 mL Ni-NTA Superflow resin (Qiagen).
2. Wash the resin with 20 column volumes (CV) of wash buffer followed by 20 CV of high salt buffer and finally 20 CV of wash buffer.
3. Elute protein as a single fraction using 3 CV of elution buffer.
4. Keep aliquots of every washing/elution step including lysate and flow through (FT).
5. Monitor the presence of AMPfret by running an SDS-PAGE gel (see 3.4.3).

#### 3.4.3. SDS-PAGE of chromatography fractions

1. Dilute 20 µL of each fraction into blue loading buffer to a final concentration of 1x loading buffer and boil the samples for 1 min at 95 °C.
2. Load 5 µL of the lysate and FT fractions and 20 µL of others in the wells and 5 µL of Precision Plus Protein Dual Color Standards.
3. Run a precast 4-20% (tris-glycine TGX) gel at 200 V (see Section 3.7.).
4. At the end of the migration, disassemble the unit, remove the gel.
5. Stain the gel for 15 min in InstantBlue staining buffer, or similar, and validate the presence of all 3 subunits of of AMPfret. (α2-mseCFPΔ11: 88 kDa, β2-cp173Venus: 57 kDa and γ1: 37 kDa; see for an example SEC fractions in **Figure 2a**).

**Figure 2.**
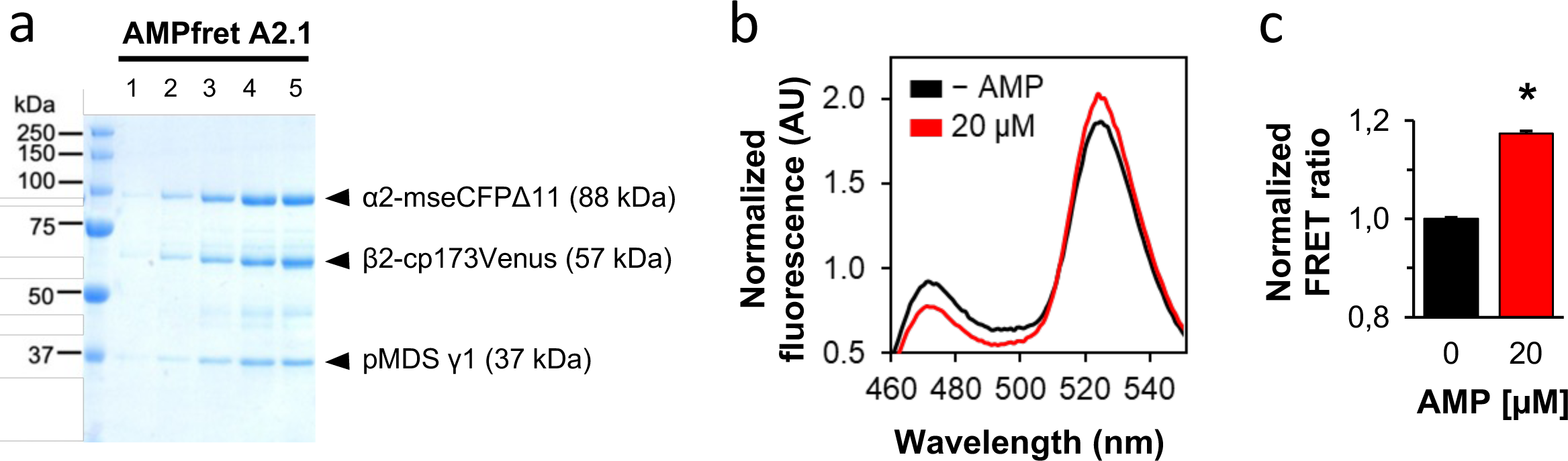

#### 3.4.4. Heparin affinity chromatography

1. Apply AMPfret protein complex eluted from IMAC onto a 5 mL Heparin column using an FPLC Akta Purifier equipped with a super-loop or equivalent.
2. Wash the column with 10 CV of Buffer A and apply a continuous gradient of Buffer B over 20 CV collecting 2 mL fractions and recording the absorbance at 280 nm.
3. Run the fractions showing absorbance at 280 nm onto an SDS-PAGE gel as described in 3.4.3 (20 µL/well) to identify where AMPfret is eluted (this will be at a point on the gradient approximately corresponding to 250 mM NaCl).

#### 3.4.5. Size exclusion chromatography (SEC)

1. Concentrate the protein complex by applying Heparin eluted fractions containing AMPfret to Amicon Ultra-4 or −15 Centrifugal Filter Unit(s) (50 kDa) pre-equilibrated with SEC buffer.
2. Centrifuge at 4,000 x g for 10 min at 4 °C in a cooled table top centrifuge. Resuspend the protein solution by pipetting up and down, being careful to not touch the centrifugal filter membrane.
3. Repeat centrifugation followed by resuspension until a final volume of 200 µL is reached. Stop centrifuging if precipitation occurs.
4. Transfer concentrated AMPfret into a 1.5 mL Eppendorf tube and spin it down at 16,000 x g for 10 min at 4 °C.
5. Inject the supernatant and run it at a flow rate of 0.5 mL/min on a Superose 6 10/300 column pre-equilibrated with SEC buffer and mounted on a FPLC AKTA purifier.
6. Collect 1 mL fractions and record the absorbance at 280 nm.
7. Run the fractions showing absorbance at 280 nm onto an SDS-PAGE gel as described in 3.4.3 (20 µL/well) to identify where AMPfret is eluted (**Figure 2a**).

#### 3.4.6. Storage

1. Pool the fractions containing pure AMPfret protein complex and concentrate them using an Amicon™ Ultra-4 or −15 centrifugal filter unit(s) (50 kDa) pre-equilibrated with SEC buffer.
2. Centrifuge at 4,000 x g for 10 min at 4 °C in a cooling table top centrifuge. Resuspend the protein solution by pipetting up and down being careful to not touch the centrifugal filter membrane.
3. Repeat centrifugation followed by resuspension and measure protein concentration using a Nanodrop device with SEC buffer as blank.
4. Concentrate until reaching a final concentration of 6 mg/mL measured using a Nanodrop.
5. Slowly add an equal volume of glycerol to the purified protein to reach a final concentration of 50 % (v/v). Mix well. The new concentration is 3 mg/mL.
6. Purified AMPfret in 50 % glycerol is stored at −20 °C until use for enzymatic assays or in vitro fluorometric FRET assays.

### 3.5. In vitro FRET assays using fluorimetry

All subsequent preparation steps are carried out on ice with cold buffers. The following protocol corresponds to the measurement of AMP-induced FRET changes. Tested AMP concentrations are 0 and 20 µM. Measurement are done in triplicates, with n being the total number of measurements (here: n=6).

1. Turn on the fluorimeter (PTI), enter the Felix software and select to record fluorescence emission scan. Set the excitation wavelength to 430 nm and the range for the emission spectrum from 450 to 600 nm with a step size of 1 nm and an integration time of 0.2 s.
2. Prepare and label Eppendorf tubes corresponding to every single measurement and place them on ice.
3. Prepare a mastermix of pure AMPfret in spectro buffer:

Pipette (n+1) * 147 µL spectro buffer and add (n+1) * 1 µL purified AMPfret at 3 mg/mL (15 pmol) (here for n=6: 1,029 mL spectro buffer + 7 µL purified AMPfret). Mix well by pulse vortexing.
4. Distribute 148 µL into each Eppendorf tube.

#### 3.5.1. Measurement

1. Add 2 µL of spectro buffer into the first tube containing 15 pmol AMPfret and mix briefly using the vortex (total 150 µL).
2. Transfer the 150 µL to the clean and dry Quartz cuvette.
3. Place the cuvette into the fluorimeter and acquire the fluorescent emission spectrum (**Figure 2b**).
4. Repeat step 1-3 for the 2 remaining replicates corresponding to basal AMPfret fluorescent emission spectrum.
5. Clean cuvette between uses: 3 times with water and 3 times with buffer.
6. Proceed the same way (steps 1-5), but adding 2 µL of AMP at 1.5 mM prepared in spectro buffer for the next 3 tubes (20 µM final).

#### 3.5.2. Data treatment

1. Import data into Excel.
2. Calculate the FRET ratio as being the ratio of emission maxima at 527 ± 2 nm and 476 ± 2 nm for each measurement.
3. Average together the values obtained for each of the replicates and normalize the obtained mean values using the averaged FRET ratio measured for AMPfret in absence of AMP (**Figure 2c**).

### 3.6. Cell-based FRET assays using microscopy

#### 3.6.1. Transient cell transfection

HEK 293T (cultured in DMEM containing 4.5 g/L glucose, 10 % FBS, penicillin and streptomycin) are transfected when the cells reach 60% confluency, 24 h to 48 h prior being observed under the microscope.

1. Trypsinise stock flask of HEK 293T cells and seed 0.05×10^6^ cells/well in 400 µL medium in an 8-well chamber slide.
2. When the next day, cells reach 60% confluency, continue with transfection.
3. Dilute 1.2 µg DNA of MultiMam AMPfret 2.1 (from Section 3.1.2) into a total volume of 62.5 µL OptiMEM. Mix by quick vortexing and spin down the solution.
4. Add 12.5 µL Polyfect transfection reagent to the DNA solution and mix by vortexing for 5 s.
5. Incubate the DNA/Polyfect mix for 10 min at room temperature.
6. Meanwhile, in each well, replace the 400 µL of complete medium with 150 µL of fresh complete medium.
7. Add 370 µL of complete medium to the transfection complexes (DNA/Polyfect) and mix by pipetting up and down twice.
8. Immediately transfer 55 µL of the transfection complexes solution into each well of the 8-well chamber slide. Gently swirl to ensure uniform distribution.
9. After 16-24 h gently aspirate the medium and add 400 µL complete medium onto the cells.
10. Proceed to FRET measurements with confocal (see Section 3.6.7) or epifluorescence microscopy (see Section 3.6.8).

#### 3.6.2. Cell transfection for generating stable cell lines

The cells are transfected for 24 h using Lipofectamine 3000, according to the following protocol:

1. Seed 0.2×10^6^ HEK 293T cells/well in a 6 well plate.
2. When, the next day, cells reach 80% confluency, continue with transfection.
3. Wash the cells twice with PBS 1x Ca2+/Mg2+.
4. Replace the medium with OptiMEM (2 mL per well).
5. Incubate the cells at least 1 h at 37 °C and 5 % CO2.
6. Prepare Reaction mix 1: 125 µl optiMEM medium, 4 µl lipofectamine 3000 solution.
7. Prepare Reaction mix 2: 125 µl of the optiMEM medium, 4 µl P3000 solution,4 µg large cargo AMPfret 7.01 vector (from Section 3.2).
8. Mix by pipetting the mixture to solubilize the DNA.
9. Add reaction mix 2 to reaction mix 1 (DNA-lipid complex, do not vortex) and incubate the mixture for 15 min at room temperature.
10. Add 250 µl per well of the DNA-lipid complex to the cells.
11. 24 h post transfection, wash the cells twice with PBS 1x Ca^2+^/Mg^2+^ and add fresh complete medium.

#### 3.6.3. Select and expand cells post-transfection

1. Select the cells after 48 h post-transfection with puromycin for two weeks. For HEK 293T cells, 1 µg/mL puromycin is appropriate, for other cell lines a proper kill curve should be established. Change the selection medium every 3 days.
2. After 2 weeks, select the HEK 293T cells with 100 µg/mL of Hygromycin B gold for at least another 2 weeks. Again, for other cell lines, proper kill curves should be established. Change the selection medium every 3 days.

#### 3.6.4. Identify single clones by limiting dilution and expansion

1. After the selection step, cease the treatment with antibiotics. Trypsinise and resuspend the transfected cells at a density of 10 cells/mL in a black 96-well cell culture plate. Add 100 µl per well (i.e., one cell per well) (see **Note 1**).
2. After 18-24 h, assess the number of cells and mark the wells with only one cell.
3. After the 4th day, for the wells identified in step 2, identify wells that now have colonies. Each colony is assumed to be clonal.
4. Verify the expression of fluorophores by the clones using epifluorescence microscopy (**Figure 3a**). Only the clones expressing CFP, Venus and iRFP 720 are taken forward; the other clones are eliminated (BFP positive cells) (see **Notes 2 and 3**). AMPfret sensor expression can also be verified by immunoblotting, since subunit size differs from endogenous AMPK (see Section 3.7. and **Figure 3b**).

**Figure 3.**
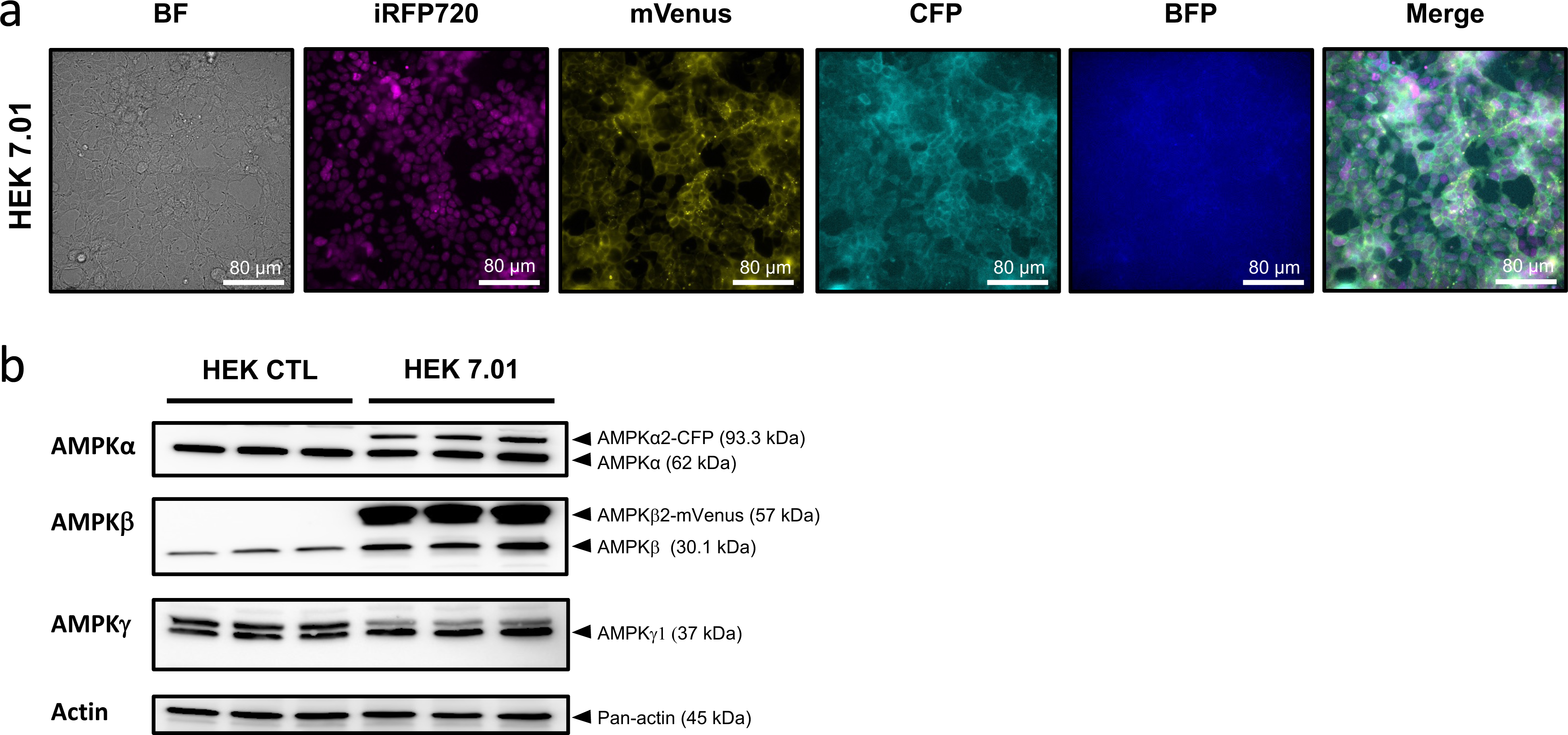
5. Once wells with CFP/Venus/iRFP 720-expressing clonal colonies have been identified, check the progress of the cells weekly until confluent.

#### 3.6.5. Transfer clones and assess expression

1. Expand the selected single-colony wells in the 96-well cell culture plate by transferring them to a 12-well cell culture plate (for the transfer, detach the cells using EDTA-trypsin, transfer the cells to the new plate.
2. When the cells are confluent in the 12-well plate, trypsinise and seed a 2-well slide for a FRET assay screen (see Sections 3.6.7. or 3.6.8.), and transfer the remainder of the cells from the 12-well plate well to a 6-well cell culture plate.
3. Keep clones that have a change in the FRET ratio following treatment and discard the others.

#### 3.6.6. Expand and freeze down high expressing clones

1. Once AMPfret 7.01 expression and FRET responses have been verified, transfer the clones to T75 flasks. Most cell clones grow fast and reach a high cell density within 2 weeks.
2. Once expanded, freeze the clones using an appropriate freezing medium lacking the selection antibiotic. A common freezing medium is 10 % dimethyl sulfoxide (DMSO) plus the appropriate complete medium for each cell line (see **Notes 4 and 5**).
3. To confirm whether the stable cell line correctly expresses and activates the heterotrimeric AMPfret complex, HEK 7.01 cells treated or not with e.g. 3 µM of FCCP are analyzed for the presence and phosphorylation of the AMPKα subunit and the phosphorylation of the AMPK substrate ACC using SDS-PAGE and immunoblotting as detailed in Section 3.7. (**Figure 4a**).

**Figure 4.**
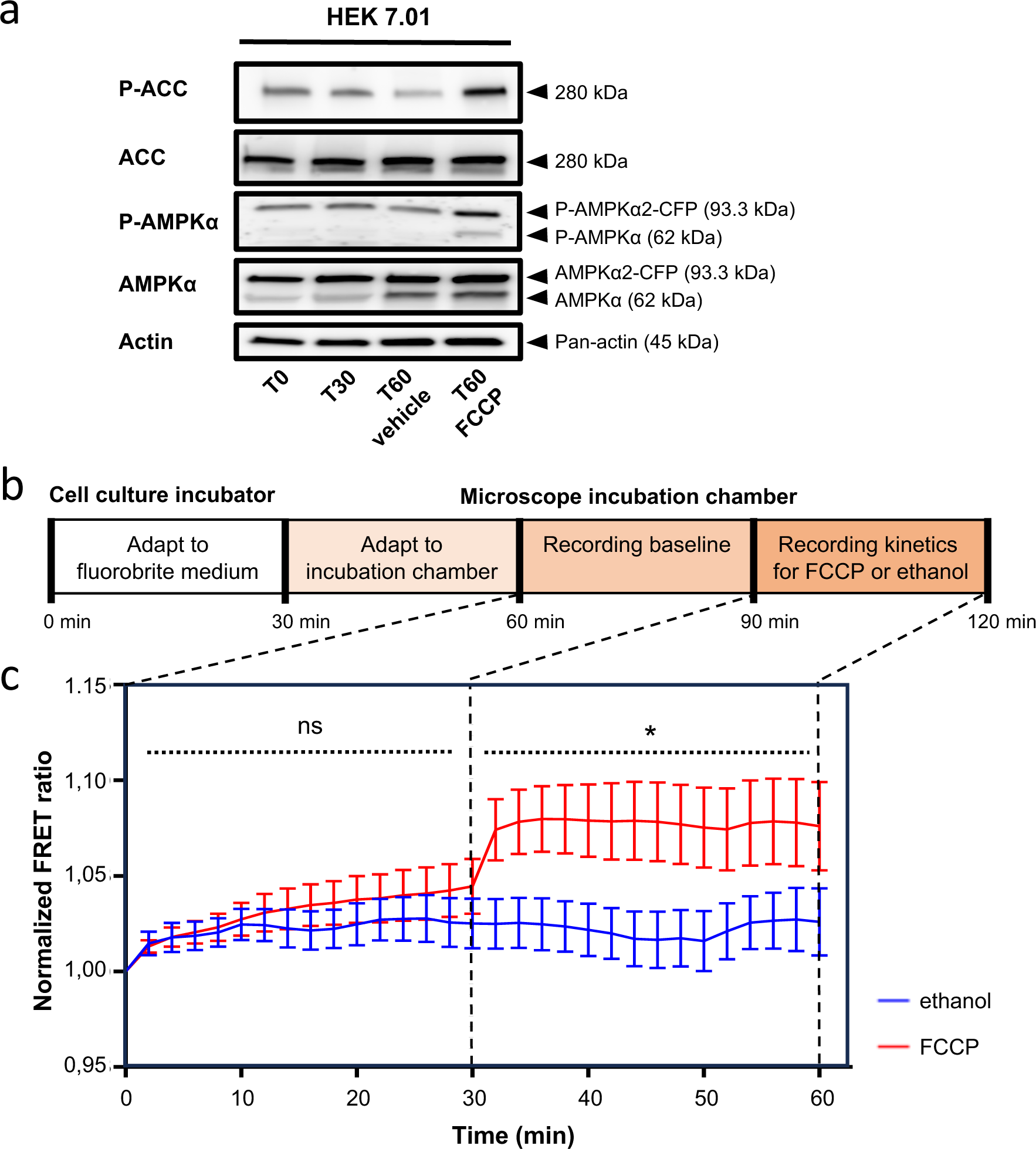

#### 3.6.7. FRET assays using confocal microscopy

The following protocol describes how to monitor AICAR-induced allosteric activation of AMPK. It can be adapted for 2-deoxyglucose by replacing the medium under the microscope with 1 g/L glucose (instead of 4,5 g/L glucose) DMEM + 10 % FBS containing 3.3 g/L 2-deoxyglucose (2-DG) (**Figure 5**).

**Figure 5.**
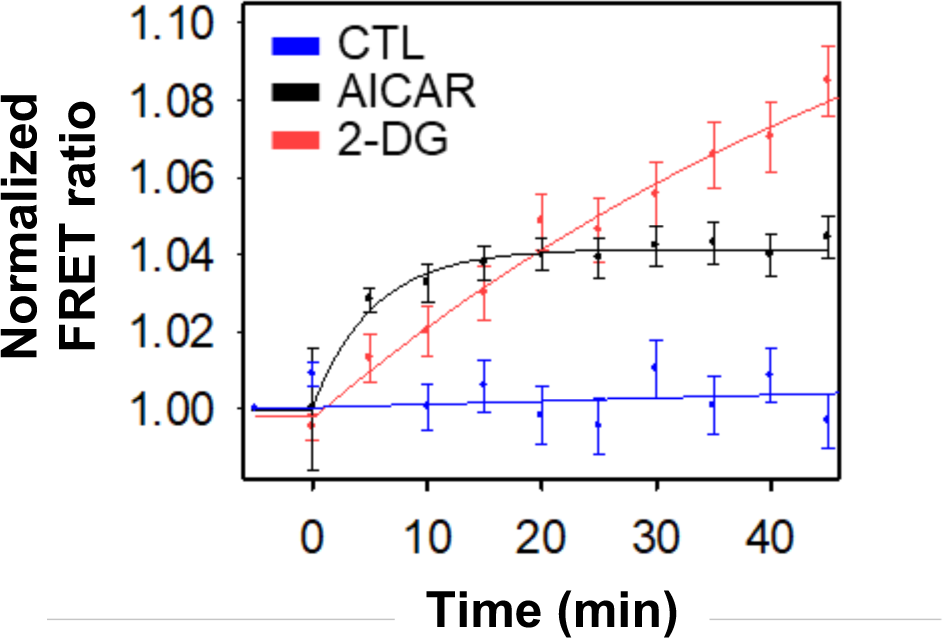

1. Place the 8-well chamber slide on the slide holder of the microscope inside the incubation chamber at 37 °C, 5 % CO_2_ and 80 % humidity (see **Note 6**). Let the cells stand there for 30 min.
2. Enter the LAS software and set the fluorescence recording settings. Excitation is done using a 458 nm argon laser (power < 20 %) and emitted fluorescences are observed through the 478 ± 5 nm and 530 ± 5 nm channels for CFP and Venus, respectively (see **Note 7**).
3. Pick a field into which several cells are visible and showing fluorescence in both CFP and Venus channels (see **Note 8**).
4. Take a picture for both channels to set the acquisition parameters (HyD or PMT gain, line/frame averaging) (see **Note 9**)
5. For each well, select a field in which several cells are visible and showing CFP- and Venus-fluorescence and retain them using the “Mark and Find” option.
6. Turn on the autofocus system.
7. Set the microscope to take a picture every 5 min over 60 min at all selected positions.
8. Run the experiment.
9. After 10 min (2 sets of pictures acquired), remove the lid of the chamber, remove 200 µL of medium, using a pipette and replace with 200 µL of complete medium containing 2 mM AICAR, prewarmed at 37 °C (1 mM final concentration in the well). Quickly repeat this for all wells of the chamber – for control wells, replace the medium with complete medium only.
10. Once all the images are recorded, save and export the experiment.
11. Import the experiment into FIJI software (see **Note 10**) and generate hyperstacks, gathering all pictures recorded over time for a given single field (i.e. one hyperstack per well with all pictures corresponding to every time point as tiles). For each hyperstack, mark every monitored cell as a region of interest. Extract CFP and Venus fluorescence intensity values at each time point for every cell.
12. Using Excel, FRET ratios are calculated for every single cell, at every time point, by dividing emissions (*em*) of Venus fluorescence by CFP fluorescence when exciting (*ex*) CFP as follows (see **Note 11**):

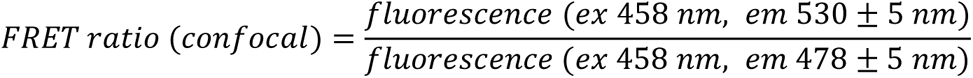
13. Results are normalized to the beginning of the experiment (0 min). Single-cell FRET ratios (for an example see **Figure 6a**) are averaged for every time point and plotted over time (i.e. data points: mean ± SEM; for an example see **Figure 6b**). See also **Note 12**.

**Figure 6.**
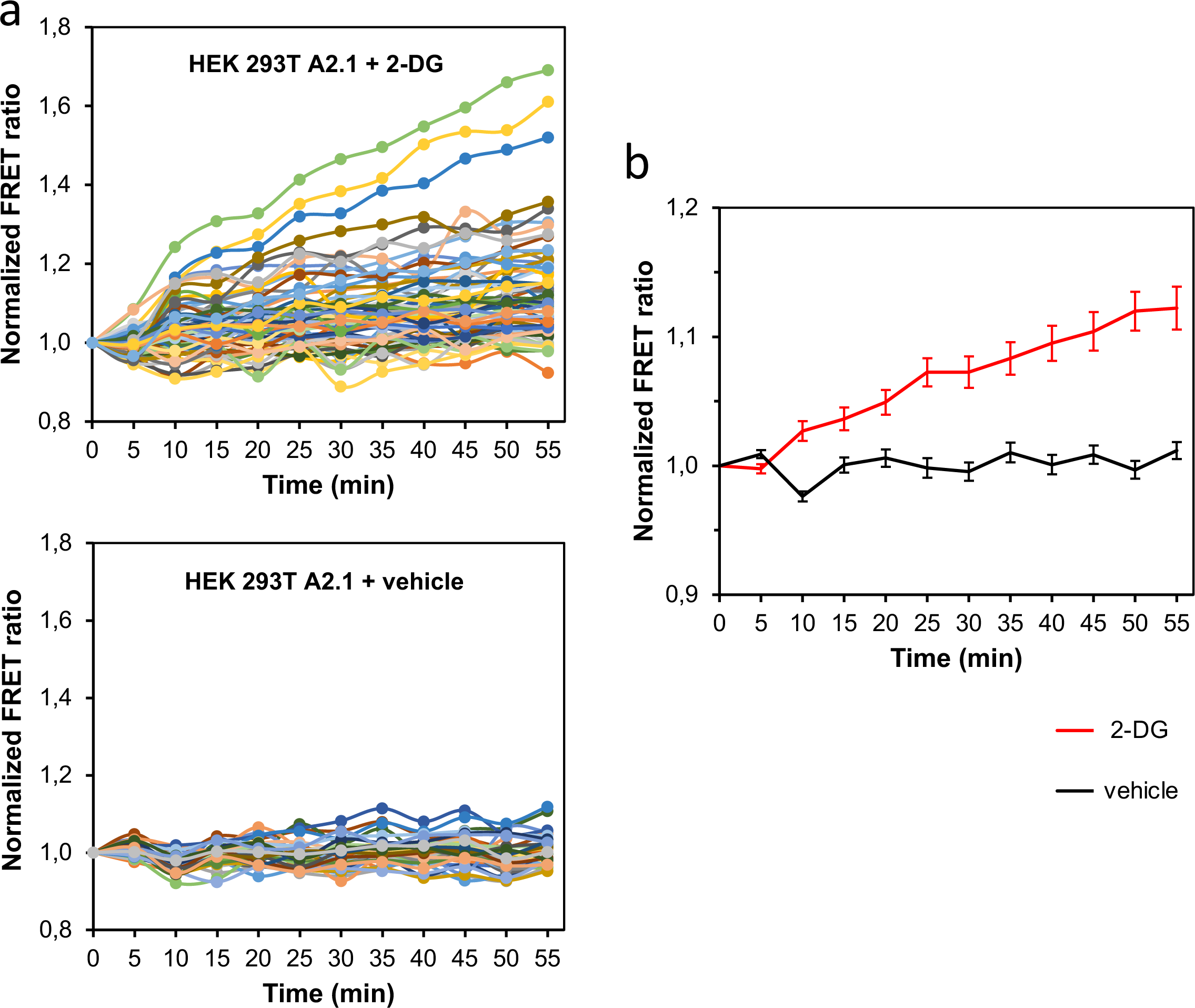

#### 3.6.8. FRET assays using epifluorescence microscopy

This assay detects FRET signal in the presence of treatments that decrease ATP/ADP and ATP/AMP ratios (e.g. AICAR, hypoxia, FCCP or oligomycin) (see **Note 1**). A typical experiment is shown in **Figure 4b**.

1. Seed 300 000 cells (stable HEK 7.01 cell line) per well in an IBIDI 2-well slide.
2. Add 1 mL of complete DMEM containing 4.5 g/L glucose, 10 % FBS, penicillin and streptomycin.
3. Allow cells to grow to reach confluence for 24-48 h.
4. Before the FRET experiments, replace the complete medium with 1 mL fluoroBrite medium supplemented with 10 % FBS, 1 mM L-glutamine and 1% Penicillin/Streptomycin (see **Note 2**).
5. Keep the cells in the incubator for 30 min at 37 °C and 5 % CO_2_ (to adapt to the new medium).
6. Put the 2-well slide in the microscope incubation chamber at 37 °C, 80 % humidity, 18 % O_2_ and 5 % CO_2_ and allow to equilibrate for 30 min.
7. Enter the FRET SE module (LAS X software, Leica).
8. Choose a field and check the microscope settings for fluorescence. The cells are excited by LEDs and fluorescence is observed with the following excitation (*ex*) and emission (*em*) wavelengths: CFP (cyan) *ex*: 440 nm, *em*: 460 nm; Venus (yellow) *ex*: 510 nm, *em*: 535 nm; iRFP (magenta) *ex*: 635 nm, *em*: 720 nm; BFP (blue) *ex*: 390 nm, *em*: 535 nm. To measure a FRET signal, you should excite CFP and observe an emission increase of Venus when AMPK undergoes an activating conformational change.
9. Activate the Adaptive Focus Control (AFC) option on the microscope so that the microscope remains focused on the cells throughout the duration of the experiment.
10. Take a capture image to configure the acquisition settings.
11. Set the microscope to acquire every 2 min and for a duration of 60 min (see **Note 3**).
12. Run the experiment.
13. After 30 min, add 3 µl of FCCP (stock solution 1 mM prepared with absolute ethanol; FCCP at 3 µM final concentration, mix by gentle pipetting) or for the control add 3 µl of absolute ethanol (vehicle). To validate the effect of FCCP, cells are cultured in triplicate under identical conditions as in the FRET assay and analyzed at 0 min and 60 min of treatment by immunoblotting for AMPK activation (**Figure 4a**).
14. Save and export the raw data.
15. The FRET ratio is calculated by dividing emission (*em*) of Venus fluorescence by CFP fluorescence when exciting (*ex*) CFP and data processed as in Section 3.6.7. (see **Note 11**):

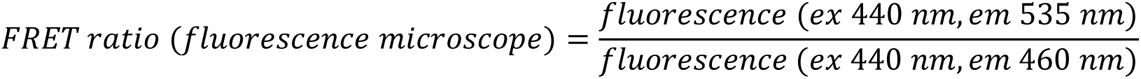
16. Use GraphPad prism8 to plot the curves. Data represent the average of the FRET ratios ± SD. Statistical tests with a 2-tailed P values ≤ 0.05 were considered significant (**Figure 4c**).

### 3.7. SDS-PAGE and immunoblotting

#### 3.7.1. Protein preparation and electrophoresis

1. Collect 2×10^6^ cells by trypsinisation and spinning in an Eppendorf tube, wash the cells with PBS Ca^2+^/ Mg^2+^ and discard the supernatant. To the pellet, add 100 µl of RIPA buffer supplemented with 1x protease inhibitor and 1x phosphatase inhibitor.
2. Incubate the homogenates on ice for 30 min and vortex samples every 5 min.
3. Centrifuge the homogenates at 15,000 × g for 30 min at 4 °C.
4. Evaluate the protein content of the samples with the smart micro-BCA protein assay kit, according to the manufacturer’s instructions using a plate reader.
5. Dilute 40 µg of proteins in blue loading buffer and boil the samples for 5 min at 95 °C.
6. Store the remaining supernatants at −80 °C until further use.
7. Assemble the gel (precast polyacrylamide gel 4-20 %) in the gel running apparatus and add the appropriate amount of running buffer (1 L). Carefully remove the comb and gently rinse the wells by injecting running buffer with a pipette
8. Load an equal amount of each sample (40 μg) in the wells and 10 μL of Precision Plus Protein Dual Color Standards.
9. Run the gel at 200 V.
10. At the end of the migration, disassemble the unit, remove the gel.

#### 3.7.2. Protein transfer and immunodetection

1. Assemble the transfer sandwich and carefully remove any air bubbles between the membrane and the gel.
2. Transfer the resolved proteins onto a 0.2 µm nitrocellulose membrane using the turbo-mixed molecular weight setting on the Trans-Blot Turbo Transfer System (Bio-Rad) or a wet-based transfer method (see **Note 4**).
3. Immerse membranes in blocking buffer for 1 h at room temperature with shaking.
4. Dilute 10 µL of anti-α-AMPK, anti-β-AMPK, anti-γ-AMPK, anti-P-T172-AMPKα, anti-ACC, anti-P-S79-ACC and anti-pan-actin antibodies in 10 mL of blocking buffer, and incubate blots overnight at 4 °C with shaking (see **Note 5**).
5. Wash membranes three times 5 min at room temperature with TBST with shaking.
6. Incubate membranes with the appropriate horseradish peroxidase-conjugated antibody for 1 h at room temperature with shaking.
7. Wash membranes three times 5 min at room temperature with TBST with shaking.
8. Incubate membranes with ECL Select Western Blotting Substrate Kit.
9. Reveal membranes with an appropriate luminescence imaging system (e.g. ImageQuant LAS 4000).

## 4. Notes

1. Alternatively, you can dilute the cells to obtain 0.8 cells / well in a 96 well plate.
2. At this stage you will obtain a mix of cells which express the fluorophores in a heterogeneous way (i.e. cells that express Venus, iRFP but not CFP, or cells that express CFP iRFP but not Venus).
3. The integration strategy is designed such that expression of H2B iRFP can only occur when the whole construct is *bona fide* integrated at the hACTB locus. This is due to the fact that the H2B iRFP is a promoterless cassette, whose expression is only restored after correct knock-in in the ACTB locus. Despite correct integration of the whole transgene cassette, most of the cells will undergo stochastic inactivation of the exogenous promoters driving alpha/beta and gamma expression. By contrast, H2B iRFP is never silenced, because its expression depends on the endogenous ACTB promoter after gene editing. Conversely, mTagBFP expression, which is placed outside the HITI2c-payload in the all-in-one construct, serves as a transfection marker and it is not meant to be integrated at the ACTB locus (**Figure 1b**). Consequently, mTagBFP expression is progressively lost after transfection and, therefore, clones exhibiting prolonged expression of mTagBFP indicate random plasmid backbone integration somewhere else in the genome. While this would not constitute a problem for any other large cargo integration approach, mTagBFP clones have to be removed due to the spectral overlap with CFP for downstream experiments (e.g. FRET microscopy or flow-cytometry). MultiMate HITI-2c H2BiRFP PH was generated by replacing the mCherry sequence in MultiMate HITI-2c 4K-CH [20] with H2BiRFP720 through Gibson assembly.
4. Before freezing, check the cells for contamination with mycoplasma, using e.g. the Lonza MycoAlert test, according to the manufacturer’s instructions together with a plate reader (e.g. CLARIOstar, BMG Labtech).
5. It is critical to follow the passage number of stable cell lines since the stability of clonal cell lines might vary. Some clones may lose expression after several passages. Freeze down samples from early passage to prolong their use after thawing.
6. One can use a small piece of double-sided tape to fix the chamber to the slide holder. This will avoid any drifting when opening the chamber for medium exchange.
7. Picture can be captured using the following parameters: 1024 x 1024 pixels resolution, 400 Hz scanning frequency and 4 lines averaging.
8. Since transient transfection yields a cell population with highly inhomogeneous expression of AMPfret, avoid picking cells that show very high or low Venus and CFP fluorescence, or dot-like bright particles.
9. Make sure Venus fluorescence intensity is not showing any saturated pixels using the saturation LUT.
10. Optional: one can apply a background subtraction simultaneously to all pictures using the rolling ball option set to a radius of 200 pixels.
11. This is the simplest calculation of FRET ratio, considering only the two emission channels without correction for excitation and emission crosstalk (sensitized emission FRET imaging). Nevertheless, it allows comparison of relative changes between two situations without requiring control sensors and extensive image processing. In case the setup does not allow to assume a constant spectral bleed-through between FRET channels, tools are available to normalize FRET pictures, e.g. by using an ImageJ plug-in [21].
12. Be aware that transient transfection, depending on cells and protocols used, can diminish the available dynamic FRET ratio range.
13. We show an assay with 3 µM of FCCP for 30 min in **Figure 4c**.
14. You can use also DMEM with no phenol red, containing 4.5 g/L glucose, 10 % FBS, penicillin and streptomycin.
15. Do not acquire data with a higher frequency than 1 picture every 2 min to avoid photobleaching.
16. The Trans-Blot Turbo Transfer System is based on the transfer of proteins using a semi-dry approach. Using the turbo setting, a complete transfer can be conducted in 7 min. Alternative wet-based transfer methods can also be used.
17. For immunoblots, α2-AMPK, β2-AMPK and γ1-AMPK are best probed on separate blots (i.e. one gel per antibody), while total α -AMPK/total ACC/actin or P-AMPK/P-ACC/actin can be probed on a single blot (cut in three appropriate pieces; i.e. one gel for 3 antibodies).

## Acknowledgements

The authors acknowledge Cécile Cottet-Rouselle, Frédéric Lamarche and Malgorzata Tokarska-Schlattner for support at LBFA. This work was funded by the French National Research Agency with the Investissements d’Avenir program (ANR-15-IDEX02) and the project betaFRET (ANR-21-CE18-0060), the Grenoble Tech Transfer Agency (SATT) Linksium, and the European Commission (EC) Framework Programme FP 7 (KBBE-2013-613879, SynSignal) in an early phase. Further support came from the SFR BEeSy, a former federal research structure at the University Grenoble Alpes, and BrisSynBio, a BBSRC/EPSRC Research Centre for synthetic biology at the University of Bristol (BB/L01386X/1).

